# Breast cancer radiogenomics analysis via computational perturbation on AI-driven multi-omics guided image synthesis

**DOI:** 10.64898/2026.01.28.702436

**Authors:** Xingying Zhang, Qian Liu

## Abstract

Radiogenomics offers a promising non-invasive approach for characterizing breast cancer (BC), yet its progress is often limited by the scarcity of cohorts containing matched imaging and multi-omics data. Recent advances in generative AI have enabled the synthesis of imaging phenotypes from genomic features, but prior work has focused on the combined influence of all genomic signals rather than isolating the effects of specific biological pathways. In this study, we introduce a perturbation-based radiogenomic framework that integrates multi-omics meta-genes with a conditional generative adversarial network (cGAN) to examine how pathway-level alterations influence synthetic BC MRI phenotypes. Seventeen meta-genes derived from Bayesian Tensor Factorization were perturbed at three levels (overexpression, base case, knockout) and synthetic DCE-MRI volumes were generated for each condition. Radiomic features were extracted using MedSAM-guided segmentation and PyRadiomics, followed by statistical evaluation using one-way ANOVA and Tukey post-hoc testing. Among the 17 pathways analyzed, only two meta-genes, representing cell cycle regulation and steroid hormone biosynthesis, produced significant and biologically interpretable changes in tumor size, heterogeneity, and textural patterns. These findings show that computational perturbation can uncover pathway-specific imaging signatures and offer mechanistic insights that complement traditional radiogenomics and explainable AI approaches. This work demonstrates the potential of perturbation-driven generative models to advance precision imaging genomics in BC.

## I. Introduction

Early detection and biomarker identification are always priorities of breast cancer (BC) management. However, BC heterogeneity, driven by genetic, molecular, and cellular variations within and between tumors, presents significant challenges [1]. Previous studies have employed multi-omics data, including copy number variation (CNV), DNA methylation, and mRNA expression, to subtype BC with significantly different prognoses [2], [3]. Oue lab has analyzed protein-protein interaction data for BC, creating a hierarchical map of BC protein communities, which led to the identification of several biomarkers with implications for mutations, survival, and targeted therapies [4]. These efforts have collectively contributed to a deeper understanding of BC heterogeneity and have provided new insights into BC personalized medicine. However, these genomics-based methods have limitations— they are relatively invasive, expensive, and cannot account for tumor morphology. Radiogenomics offers a promising alternative, combining imaging and genomic data to improve BC diagnosis in a non-invasive, cost-effective manner [5]. Previous studies have demonstrated that radiogenomic analysis can reveal genetic details at the voxel level within a heterogeneous tumor, aiding personalized treatment strategies [6]. We have conducted several radiogenomics studies for both BC and neurodegenerative diseases, resulting in a series of publications [7], [8], [9], [10]. A common challenge in BC radiogenomics lies in the need for medical imaging, genomic, and clinical outcome data from the same cohort—a requirement that is often difficult to fulfill due to the logistical and resource-intensive nature of data collection. Addressing this limitation will be key to advancing the field and unlocking the full potential of radiogenomics in clinical applications.

Recent advancements in deep generative models, such as GPT, have made it possible to synthesize medical images based on genomic information. There are several generative AI models, such as AutoEncoder (AE), Variational AutoEncoder (VAE), Transformer, Generative Adversarial Networks (GAN), diffusion model and others, which have shown impressive results in generating medical images for organs like the brain, liver, and lungs [11], [12], [13], [14], [15]. However, studies focusing on BC remain limited. Our research group is among the first to generate BC images using generative models. In our first study, we used Least Absolute Shrinkage and Selection Operator (LASSO)-based multi-variable regression to predict image features from multi-omics genomic data and successfully identified significant survival biomarkers for BC [9]. Later, we applied Conditional GAN (cGAN) to generate 3D MRI volumes from multi-omics features, enabling accurate predictions of BC subtypes and mutations [16]. More recently, we developed an advanced Conditional Probabilistic Diffusion Model (CPDM) to generate BC MRI images. The generated MRIs performed very well in predicting TP53 mutation status, ER status, BC subtypes, and survival rates [7]. However, these generative models conditioned on the “omics” level gene information, primarily focused on the combined effects of the entire genome on imaging features, which provides a broad understanding but lacks specificity. They did not delve into the individual contributions of specific genes/gene sets, leaving a gap in understanding how particular genetic/pathway alterations directly influence imaging characteristics. By not isolating the effects of gene sets, these studies missed an opportunity to identify precise genetic biomarkers that could enhance personalized treatment strategies and improve our understanding of tumor heterogeneity. Current approaches often rely on explainable AI (xAI) tools to evaluate the gradient or importance of each genetic input in the model’s decision-making process [10]. However, these methods focus on explaining the model’s predictions rather than specifically isolating and analyzing each meta-gene’s direct effect on imaging changes.

Perturbation-based methods, such as those used in Dependency Map (DepMap) [17] and Connectivity Map (cMap) [18], offer a promising strategy for addressing the gap in understanding the effects of individual meta-genes on imaging characteristics. These approaches enable the isolation and study of specific meta-gene effects on phenotypes, providing a level of granularity that is currently missing in generative models conditioned on “omics”-level gene information. DepMap, for instance, employs large-scale CRISPR screening to systematically map genetic dependencies in cancer, allowing researchers to identify which genes are critical for tumor survival and growth [17]. Similarly, cMap connects drug treatments to changes in gene expression, offering insights into how specific genetic alterations interact with therapeutic compounds [18]. The success of these resources highlights the power of perturbation approaches to uncover critical insights into the molecular mechanisms underlying disease phenotypes. Applying such perturbation-based strategies to radiogenomics could fill the current gap by enabling researchers to assess how individual genetic alterations drive specific imaging features. This would provide a more precise understanding of tumor heterogeneity and enable the discovery of actionable genetic biomarkers.

This study is to bridge the gap in understanding the specific contributions of individual gene sets to imaging phenotypes in BC through the application of AI-driven computational perturbations. Our hypothesis is that by systematically assessing the effects of genetic alterations on image synthesis, we can identify novel biomarkers that provide a deeper understanding of tumor biology and enable more precise stratification of BC heterogeneity. Such an approach addresses the current limitations of xAI methods. This research aims to extend such methodologies into the BC radiogenomics domain.

## II. Materials and Methods

### A. Dateset

Multi-omic data consisted of three varying types of genomic data (CNV score, gene expression, DNA methylation) and was retrieved from the Breast Invasive Carcinoma (BRCA) project in The Cancer Genome Atlas (TCGA) platform [19]. After matching, there were 754 patients with all three omics data and a multi-omics tensor was constructed from these three sources. The tensor was then decomposed using the Bayesian Tensor Factorization (BTF) algorithm to generate 17 latent features (also called meta-genes) for each patient. Please refer to our previous publications for more details about the BTF multi-omics tensor extraction of the 17 multi-omics meta-genes [2], [3]. In previous study, we also did gene set enrichment analysis (GSEA) to check which pathway each meta-gene is likely representing. This increased the interpretability of the meta-genes, which helped a lot in the discussion of our perturbation analysis in this study.

Dynamic contrast-enhanced MRI (DCE-MRI) scans for BC patients were retrieved from The Cancer Imaging Archive (TCIA) [20]. Within the TCGA-BRCA cohort, 754 patients possessed complete multi-omics profiles; however, only 61 of these individuals had corresponding DCE-MRI data available. This stark imbalance exemplifies the unpaired-data challenge common to BC repositories: extensive genomic data exist, but matched imaging data remain scarce. The raw DCE-MRI scans were provided in the Digital Imaging and Communications in Medicine (DICOM) format, which contains comprehensive acquisition metadata. For the purposes of this study, only the pixel-level image information was used. Across the 61 patients, 187 3D DCE-MRIs were available, reflecting the fact that each patient may undergo multiple acquisitions. These 3D volumes were captured at sequential time points separated by tens of seconds to characterize dynamic contrast uptake patterns. To ensure comparability across scans, all 3D volumes were spatially normalized to a uniform resolution of 32 × 128 × 128 voxels. The 187 scans comprised two anatomical perspectives: a side (lateral) view and a top-down (axial) view. Among them, 58 were side-view volumes and 129 were top-down volumes. Although the axial view was more prevalent, the side view was selected for model development because it provides a more intuitive depiction of breast morphology and allows clearer visual assessment of image quality, particularly when evaluating synthetic MRI outputs. The 58 side-view DCE-MRIs were subsequently partitioned into training and testing subsets of the cGAN, containing 50 and 8 volumes respectively.

### B. Methods

The overall workflow of the study is illustrated in **Figure 1**. The 17 multi-omics meta-genes were used as conditioning variables for a cGAN, enabling the generation of synthetic BC DCE-MRI volumes based on specific multi-omics perturbations. Details regarding the design, training, and performance of the generative model can be found in our prior publication [16].

**Fig. 1.**
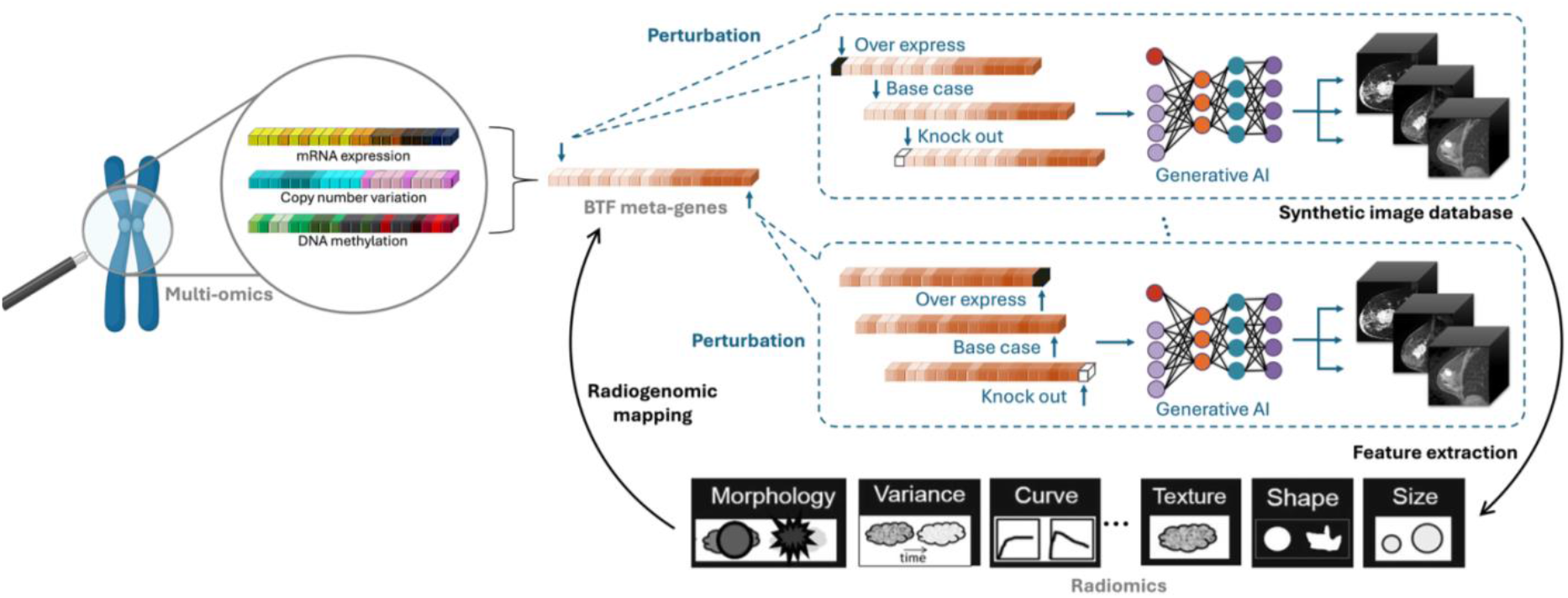
The overall workflow of this study.

To investigate how individual biological processes influence MRI-derived tumor phenotypes, each of the 17 meta-genes was perturbed at three biologically meaningful levels:

- Overexpression: The meta-gene value was increased to its maximum observed level across all patients, simulating a strong upregulation of the associated biological pathway.
- Base case: The original, unaltered meta-gene value was used as the reference condition.
- Knockout: The meta-gene value was set to zero, mimicking complete suppression of the pathway’s activity.

For each perturbation condition, we constructed a modified 17-dimensional meta-gene expression vector. This vector was then fed into the trained cGAN to generate 15 synthetic DCE-MRI volumes per condition, producing a total of 45 volumes per meta-gene and 765 volumes across all 17 meta-genes. All generated MRI volumes underwent tumor segmentation guided by MedSAM [21], which is a pretrained segmentation model for medical images. After which radiomic feature extraction was performed using PyRadiomics [22]. A set of 32 radiomic features was extracted from each volume, representing first-order intensity statistics and texture descriptors derived from Gray Level Co-occurrence Matrix (GLCM), Gray Level Dependence Matrix (GLDM), and Neighborhood Gray Tone Difference Matrix (NGTDM) feature families. These features quantify tumor size, intensity distribution, homogeneity, heterogeneity, and local structural patterns.

To evaluate whether perturbations of individual meta-genes resulted in measurable differences in radiomic signatures, we conducted a One-Way Analysis of Variance (ANOVA) for each feature–meta-gene pair. This allowed us to determine whether the overexpression, base case, and knockout groups exhibited statistically significant differences. When the ANOVA indicated significance (p < 0.05), we performed Tukey’s Honest Significant Difference (HSD) post-hoc tests to identify which pairwise comparisons (overexpression vs. base case, overexpression vs. knockout, base case vs. knockout) differed significantly.

## III. Results

Figure 2. presents a heatmap of the 17 meta-genes extracted using the BTF method. Each meta-gene represents a latent multi-omics component derived from the joint structure of CNV, DNA methylation, and gene expression data [2], [3]. The heatmap displays the expression levels of these meta-genes for the 58 patients included in this study. As detailed in **Table 1**, each meta-gene has been annotated with its most enriched biological pathway based on gene set enrichment analysis from our previous work. These pathways cover a broad spectrum of biological processes, including immune signaling (for example, chemokine signaling and cytokine receptor interaction), metabolic processes (such as steroid hormone biosynthesis and xenobiotic metabolism), and cancer-relevant mechanisms (such as cell cycle regulation and VEGF signaling). This annotation provides a biologically interpretable basis for understanding how perturbations of individual meta-genes may influence imaging phenotypes in subsequent experiments.

**TABLE I.**
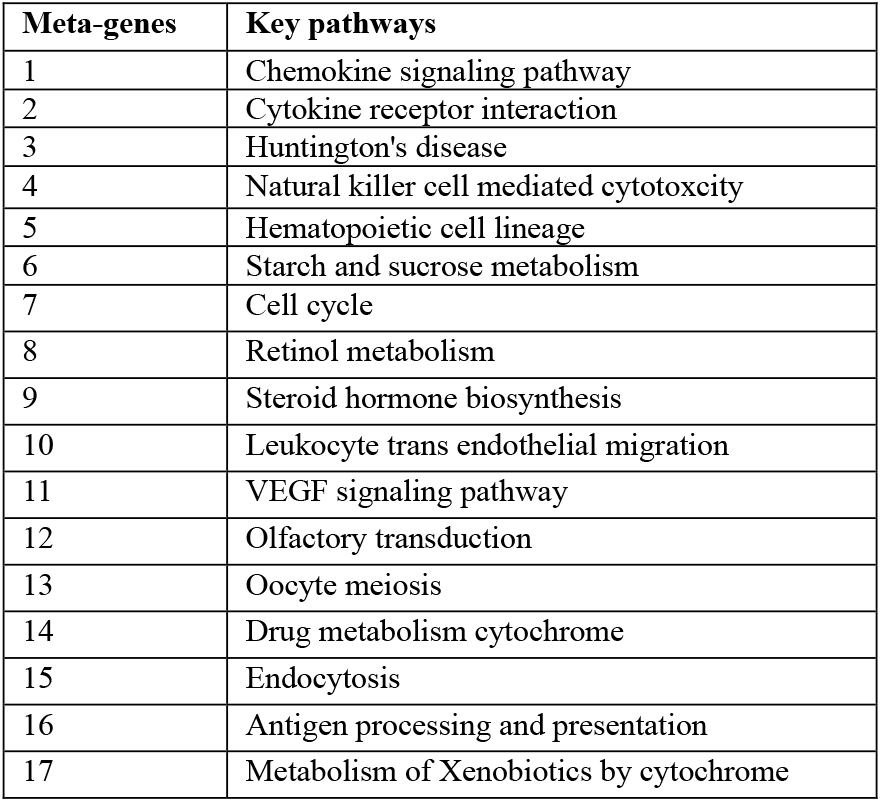
Key biological pathways for each meta-gene.

**Fig. 2.**
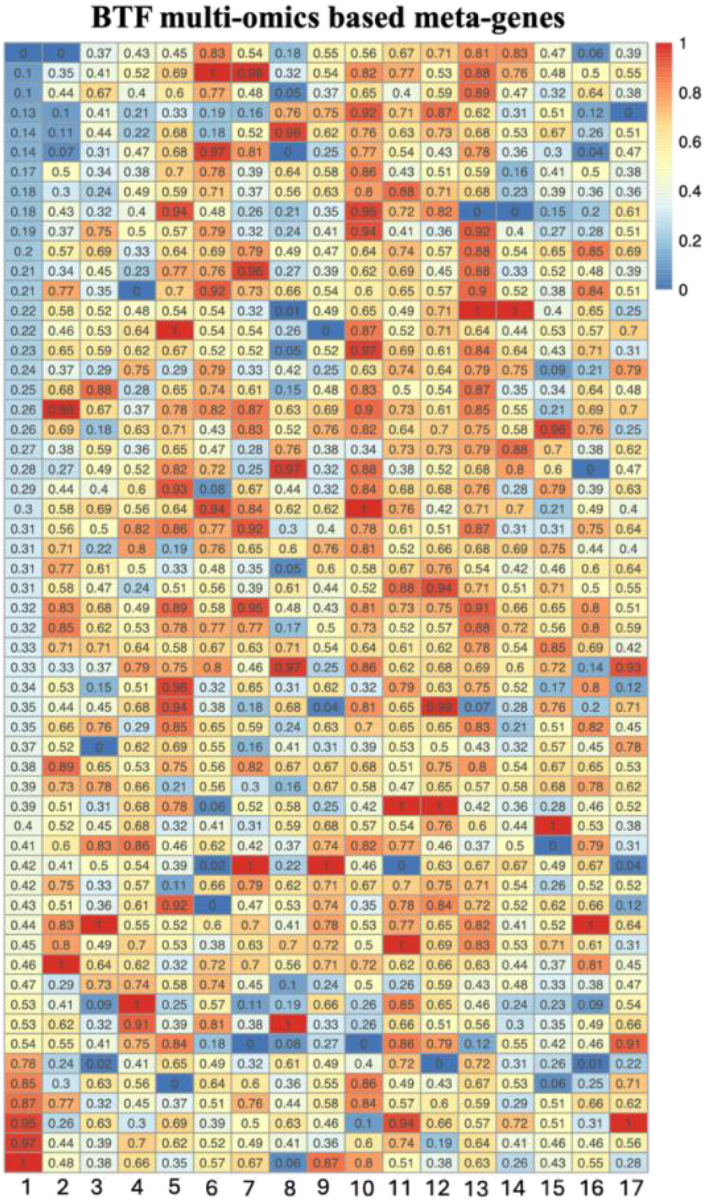
A heatmap showing the 17 BTF meta-genes extracted from multi-omics data using BTF method for 58 patients.

Figure 3. illustrates representative synthetic DCE-MRI volumes generated under three perturbation conditions (overexpression, base case, and knockout) for the two meta-genes found to have significant effects in our analysis: BTF#7 and BTF#9. Consistent with its biological role in regulating cell proliferation, perturbing the cell cycle meta-gene (BTF#7) produced visible changes in tumor morphology, including variations in apparent tumor size and boundary irregularity. In contrast, perturbation of BTF#9, associated with steroid hormone biosynthesis, resulted in more subtle but noticeable differences in texture smoothness and structural uniformity, reflecting hormone-driven alterations in tissue composition commonly observed in breast cancer. These examples demonstrate that pathway-specific multi-omics signals can manifest as distinct radiomic phenotypes in the generated MRI volumes.

**Fig. 3.**
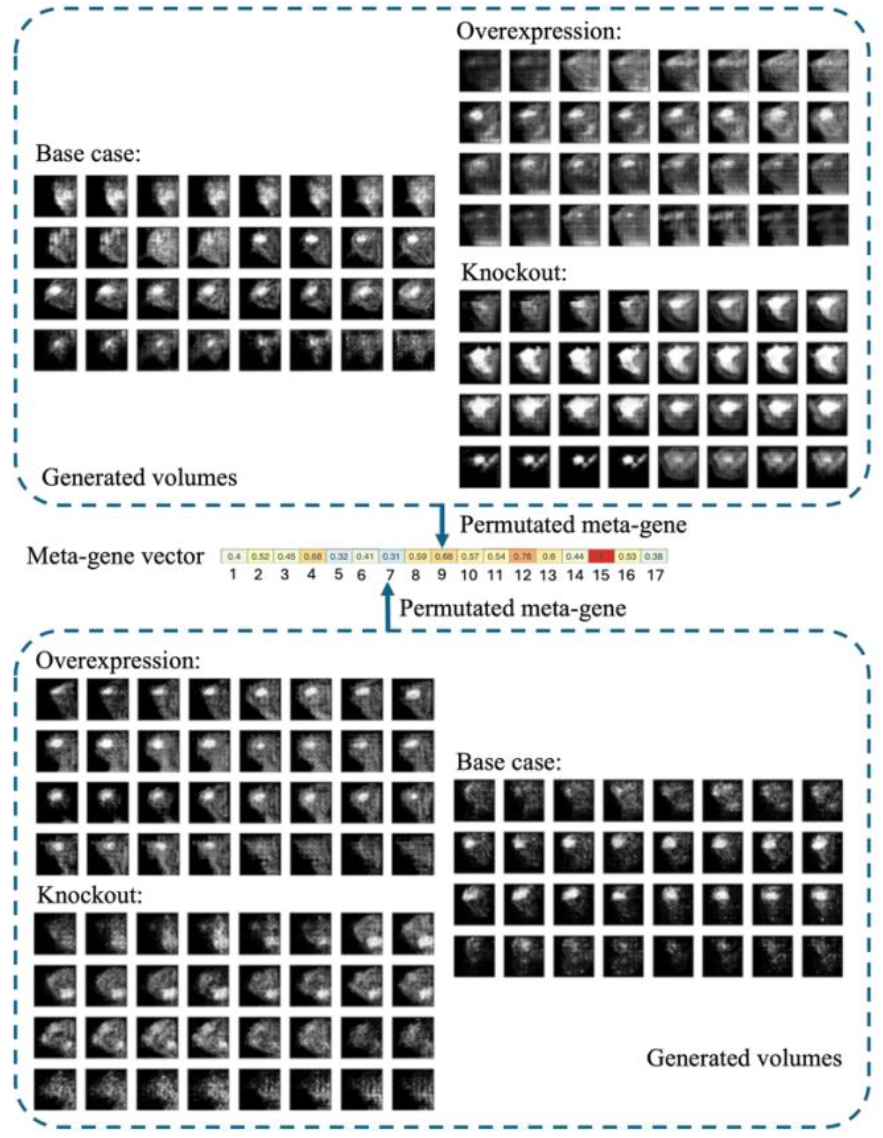
Examples of synthetic DCE-MRI volumes generated under three perturbation levels, overexpression, base case, and knockout, for the two significant BTF meta-genes (BTF#7 and BTF#9).

**Table 2** summarizes the 32 radiomic features extracted from each synthetic MRI volume, capturing a comprehensive set of descriptors related to tumor size, intensity distribution, and textural heterogeneity. These features span first-order statistics, GLCM features, GLDM features, and NGTDM features, providing a quantitative representation of the generated imaging phenotypes.

**TABLE II.**
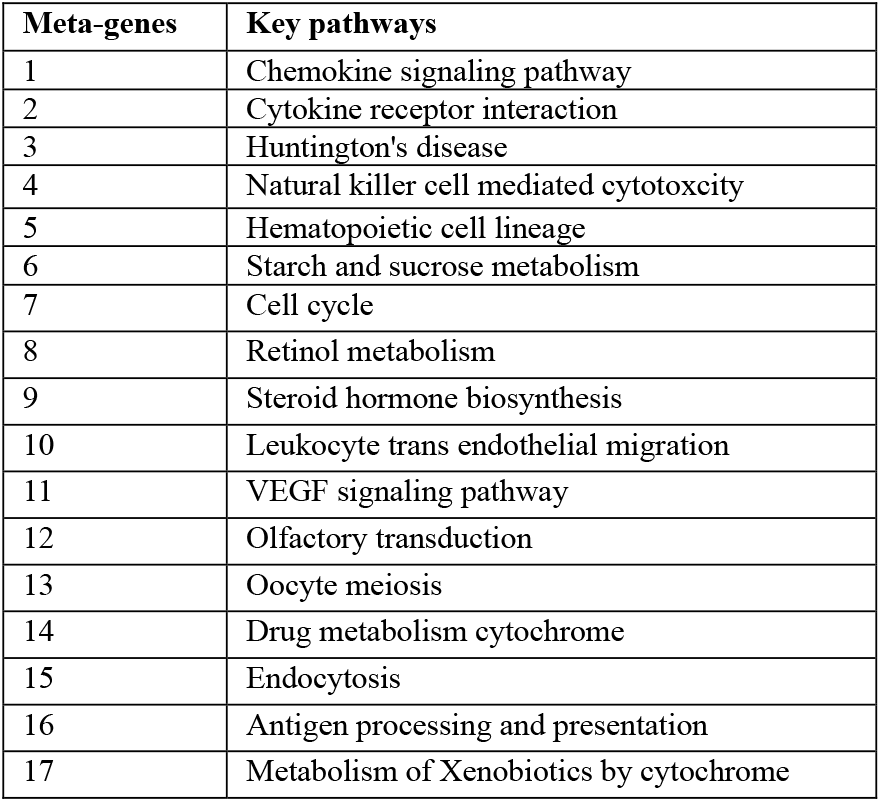
Key biological pathways for each meta-gene.

To evaluate the impact of meta-gene perturbations on MRI-derived radiomic profiles, we conducted a One-Way ANOVA followed by Tukey post-hoc comparisons. **Table 3** reports the statistically significant results. Among the 17 meta-genes analyzed, only two (BTF#7 and BTF#9) exhibited significant perturbation effects on radiomic features. For BTF#7 (cell cycle), three radiomic features showed significant differences across the perturbation levels: Area, RMS, and GLCM Contrast. Overexpression of this meta-gene resulted in larger tumor regions and increased textural variation, suggesting enhanced tumor proliferation and architectural heterogeneity, while knockout produced smaller and more homogeneous structures. For BTF#9 (steroid hormone biosynthesis), significant effects were observed in Dependence Entropy, Busyness, and Homogeneity2. These changes indicate pathway-specific influences on tissue smoothness and microstructural complexity, aligning with known effects of hormonal regulation on breast tissue organization.

**TABLE III.**
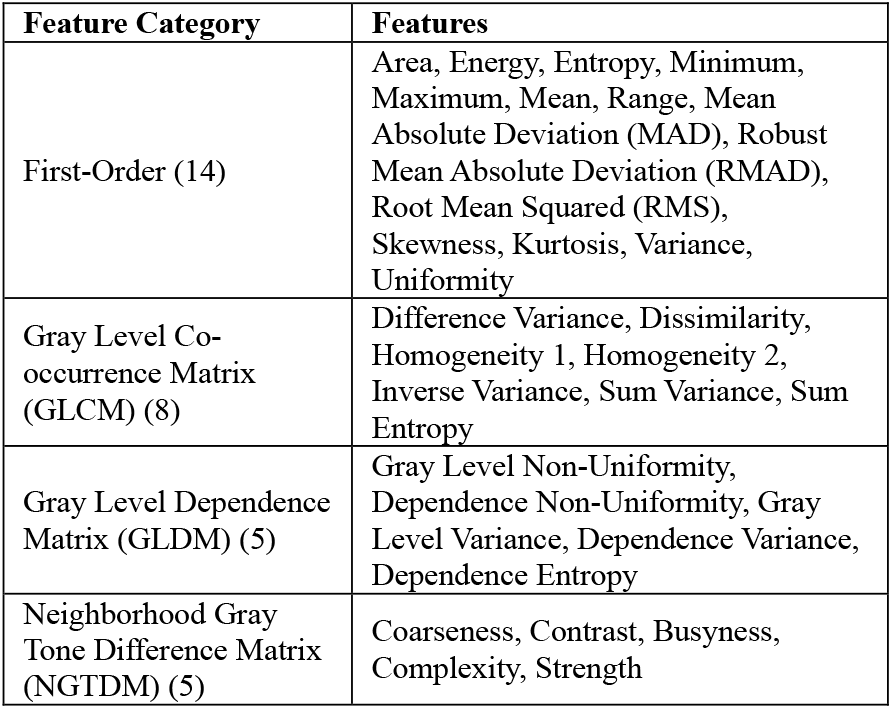
Extracted Radiomic Features Used for Tumor Phenotype Quantification.

**TABLE IV.** One-way anova and Tukey Post-hoc Tests for Perturbation Effects on Radiomic Features.

| Meta-gene | Radiomic Feature | Mean (OE) | Mean (B) | Mean (KO) | ANOVA F | ANOVA p | OE-B (p) | OE-KO (p) | B-KO (p) |
| --- | --- | --- | --- | --- | --- | --- | --- | --- | --- |
| BTF#7 | Area | 62.4 | 51.8 | 42.2 | 6.82 | 0.002** | +10.6 (0.048*) | +20.2 (0.001**) | +9.6 (0.091) |
|  | RMS | 41.3 | 38.7 | 36.9 | 4.95 | 0.010* | +2.6 (0.072) | +4.4 (0.009**) | +1.8 (0.214) |
|  | Contrast | 0.214 | 0.187 | 0.165 | 5.41 | 0.007** | +0.027 (0.031*) | +0.049 (0.004**) | +0.022 (0.067) |
| BTF#9 | Dependence Entropy | 2.87 | 2.74 | 2.52 | 8.12 | <0.001*** | +0.13 (0.044*) | +0.35 (0.0004***) | +0.22 (0.016*) |
|  | Busyness | 0.186 | 0.170 | 0.154 | 5.89 | 0.005** | +0.016 (0.083) | +0.032 (0.007**) | +0.016 (0.048*) |
|  | Homogeneity2 | 0.412 | 0.398 | 0.365 | 4.02 | 0.023* | +0.014 (0.094) | +0.047 (0.009**) | +0.033 (0.041*) |

OE: overexpress; B: base case; KO: knockout

## IV. Discussion and Conclusion

This study introduces a perturbation-based radiogenomic framework that integrates multi-omics derived meta-genes with a generative AI model to evaluate how specific biological pathways influence BC MRI phenotypes. Unlike prior radiogenomic models that characterize only the global genomic contribution to imaging, our approach isolates the effect of each meta-gene by systematically perturbing individual components of the 17-dimensional multi-omics representation. This design enables pathway-level interpretability that is not achievable using standard explainable AI methods, which typically provide importance scores but do not model how changes in specific biological signals translate into corresponding imaging phenotypes.

Our results highlight two meta-genes, BTF#7 (cell cycle) and BTF#9 (steroid hormone biosynthesis), as major contributors to synthetic MRI variation. Perturbing these pathways produced significant alterations in radiomic signatures, including changes in tumor size, heterogeneity, and textural complexity. These findings are consistent with well-established BC biology [23]. Dysregulation of the cell cycle is a fundamental driver of tumor proliferation, while steroid hormone signaling plays a central role in shaping tissue morphology and microenvironmental organization in hormone-responsive tumors [23]. The ability of the generative model to reproduce these biologically meaningful patterns suggests that multi-omics conditioned image synthesis, when combined with systematic perturbation, can capture mechanistic links between genomic alterations and imaging appearance.

The distinct radiomic patterns resulting from perturbation provide further support. Overexpression of the cell cycle meta-gene increased features such as Area, RMS, and GLCM Contrast, reflecting accelerated tumor expansion and increased architectural irregularity. In contrast, perturbations of the steroid hormone biosynthesis pathway affected Homogeneity and Dependence Entropy, indicating smoother, hormonally influenced tissue characteristics commonly observed in estrogen-driven BC. This separation of phenotypic effects demonstrates that pathway-level perturbation can reveal imaging signatures associated with specific biological processes and offers a new way to interpret tumor heterogeneity using generated images.

It is important to note that the perturbations in this study operate at the meta-gene level, where each latent component represents a biologically meaningful mixture of molecular signals summarized through tensor factorization. Although this abstraction provides stability and interpretability, it reduces genetic resolution. Perturbing individual genes, curated gene modules, or specific mutations would offer finer biological granularity and may uncover additional imaging–genomic relationships that meta-gene perturbations cannot identify. As imaging-genomic datasets continue to expand, extending this framework to more granular perturbation units represents a promising direction for future research.

In conclusion, this study demonstrates that AI-driven computational perturbation of multi-omics meta-genes provides a powerful framework for uncovering how specific biological processes shape BC imaging phenotypes. By systematically altering each meta-gene within a generative model, we identified two key pathways, cell cycle regulation and steroid hormone biosynthesis, that produce distinct and measurable effects on synthetic MRI-derived radiomic features. These results illustrate the potential of pathway-level perturbation to reveal biologically meaningful imaging signatures and enhance radiogenomic interpretability. Although our current implementation focuses on meta-gene perturbation, future work that performs perturbation at the level of individual genes or gene modules may yield even more precise insights into the molecular drivers of tumor heterogeneity and further advance precision imaging genomics.

## References

[1] G. Turashvili and E. Brogi, “Tumor heterogeneity in breast cancer,” Front Med (Lausanne), vol. 4, no. DEC, p. 322799, Dec. 2017, doi: 10.3389/FMED.2017.00227/BIBTEX.

[2] Q. Liu, B. Cheng, Y. Jin, and P. Hu, “Bayesian tensor factorization-drive breast cancer subtyping by integrating multi-omics data,” J Biomed Inform, vol. 125, p. 103958, Jan. 2022, doi: 10.1016/J.JBI.2021.103958.

[3] Q. Liu, S. Huang, Z. Zhang, T. M. Lakowski, W. Xu, and P. Hu, “Multiomics-Based Tensor Decomposition for Characterizing Breast Cancer Heterogeneity,” Machine Learning Methods for Multi-Omics Data Integration, pp. 133–150, 2024, doi: 10.1007/978-3-031-36502-7_8.

[4] X. Zhang and Q. Liu, “A Graph Neural Network Approach for Hierarchical Mapping of Breast Cancer Protein Communities,” Jun. 2024, doi: 10.21203/RS.3.RS-4478708/V1.

[5] M. A. Mazurowski, “Radiogenomics: What It Is and Why It Is Important,” Journal of the American College of Radiology, vol. 12, no. 8, pp. 862–866, 2015, doi: 10.1016/j.jacr.2015.04.019.

[6] K. Pinker, J. Chin, A. N. Melsaether, E. A. Morris, and L. Moy, “Precision medicine and radiogenomics in breast cancer: New approaches toward diagnosis and treatment,” Radiology, vol. 287, no. 3, pp. 732–747, 2018, doi: 10.1148/radiol.2018172171.

[7] L. Chen, Z. H. Huang, Y. Sun, M. Domaratzki, Q. Liu, and P. Hu, “Conditional probabilistic diffusion model driven synthetic radiogenomic applications in breast cancer,” PLoS Comput Biol, vol. 20, no. 10, p. e1012490, Oct. 2024, doi: 10.1371/JOURNAL.PCBI.1012490.

[8] Q. Liu and P. Hu, “Radiogenomic association of deep MR imaging features with genomic profiles and clinical characteristics in breast cancer,” Biomark Res, vol. 11, no. 1, pp. 1–11, 2023, doi: 10.1186/s40364-023-00455-y.

[9] Q. Liu and P. Hu, “A novel integrative computational framework for breast cancer radiogenomic biomarker discovery,” Comput Struct Biotechnol J, May 2022, doi: 10.1016/J.CSBJ.2022.05.031.

[10] Q. Liu and P. Hu, “Extendable and explainable deep learning for pan-cancer radiogenomics research,” Curr Opin Chem Biol, vol. 66, p. 102111, Feb. 2022, doi: 10.1016/J.CBPA.2021.102111.

[11] A. Vaswani et al., “Attention is all you need,” in Advances in neural information processing systems, 2017, pp. 5998–6008.

[12] R. Rombach, A. Blattmann, D. Lorenz, P. Esser, and B. Ommer, “High-Resolution Image Synthesis With Latent Diffusion Models,” in Proceedings of the IEEE/CVF conference on computer vision and pattern recognition, 2022, pp. 10684–10695. Accessed: Jan. 04, 2025. [Online]. Available: https://github.com/CompVis/latent-diffusion

[13] R. R. T. Rejusha and S. V. K. Vipin Kumar, “Artificial MRI Image Generation using Deep Convolutional GAN and its Comparison with other Augmentation Methods,” ICCISc 2021 - 2021 International Conference on Communication, Control and Information Sciences, Proceedings, Jun. 2021, doi: 10.1109/ICCISC52257.2021.9484902.

[14] M. Boulanger et al., “Deep learning methods to generate synthetic CT from MRI in radiotherapy: A literature review,” Physica Medica: European Journal of Medical Physics, vol. 89, pp. 265–281, Sep. 2021, doi: 10.1016/J.EJMP.2021.07.027.

[15] W. Li et al., “Magnetic resonance image (MRI) synthesis from brain computed tomography (CT) images based on deep learning methods for magnetic resonance (MR)-guided radiotherapy,” Quant Imaging Med Surg, vol. 10, no. 6, pp. 1223236–1221236, Jun. 2020, doi: 10.21037/QIMS-19-885.

[16] Z. H. Huang, L. Chen, Y. Sun, Q. Liu, and P. Hu, “Conditional generative adversarial network driven radiomic prediction of mutation status based on magnetic resonance imaging of breast cancer,” J Transl Med, vol. 22, no. 1, pp. 1–13, Dec. 2024, doi: 10.1186/S12967-024-05018-9/TABLES/5.

[17] A. Tsherniak et al., “Defining a Cancer Dependency Map,” Cell, vol. 170, no. 3, pp. 564–576, 2017, doi: 10.1016/j.cell.2017.06.010.Defining.

[18] A. Subramanian et al., “A Next Generation Connectivity Map: L1000 Platform and the First 1,000,000 Profiles,” Cell, vol. 171, no. 6, pp. 1437–1452.e17, Nov. 2017, doi: 10.1016/J.CELL.2017.10.049.

[19] K. Tomczak, P. Czerwińska, and M. Wiznerowicz, “The Cancer Genome Atlas (TCGA): An immeasurable source of knowledge,” 2015. doi: 10.5114/wo.2014.47136.

[20] M. Zanfardino, K. Pane, P. Mirabelli, M. Salvatore, and M. Franzese, “TCGA-TCIA impact on radiogenomics cancer research: A systematic review,” Int J Mol Sci, vol. 20, no. 23, 2019, doi: 10.3390/ijms20236033.

[21] J. Ma, Y. He, F. Li, L. Han, C. You, and B. Wang, “Segment anything in medical images,” Nature Communications 2024 15:1, vol. 15, no. 1, pp. 654–, Jan. 2024, doi: 10.1038/s41467-024-44824-z.

[22] J. J. M. Van Griethuysen et al., “Computational radiomics system to decode the radiographic phenotype,” Cancer Res, vol. 77, no. 21, pp. e104–e107, Nov. 2017, doi: 10.1158/0008-5472.CAN-17-0339.

[23] D. Hanahan and R. A. Weinberg, “Hallmarks of cancer: The next generation,” 2011. doi: 10.1016/j.cell.2011.02.013.

